# Generation of High-fidelity Blastocyst-like Structures from Porcine Expanded Pluripotent stem cell via Chemically inducing cell plasticity

**DOI:** 10.1101/2024.07.24.604931

**Authors:** Zheng Liao, Xin Qi, Shijie Yuan, Junfei Zhang, Na hai, Xin Gao, Yang Yuan, Xuguang Du, Shiqiang Zhang, Yulei Wei

**Author notes:** Correspondence to: Yulei Wei;, Shiqiang Zhang, Xuguang Du. These authors contributed equally to this work (Zheng Liao, Xin Qi, Shijie Yuan).

## Abstract

Understanding the mechanisms of blastocyst formation and implantation is crucial for advancing farm animal reproduction. However, research in this area is often hindered by the limited availability of embryos. In this study, we developed an efficient method to generate blastocyst-like structures (termed blastoids) from porcine expanded pluripotent stem cells (pEPSCs) through chemical induction with an efficiency up to 80%. These porcine blastoids closely resemble natural blastocysts in terms of morphology, cell composition, and single-cell transcriptomes. The tissue structures of the blastoids developed over time in culture, resembling the tissue morphology of gastrulation-stage embryos. This innovative approach not only provides a robust in vitro model for studying early embryogenesis in pigs but also holds the potential for improving reproductive efficiency and understanding the developmental processes of large farm animal species. The development of porcine blastoids represents a significant advancement in regenerative medicine and developmental biology.

## Introduction

The formation and implantation of blastocysts are pivotal events in mammalian reproduction, essential for understanding human reproductive health and enhancing farm animal breeding practices. However, the scarcity of embryos available for research with pigs has historically limited our ability to comprehensively study these processes. Recent advancements in stem cell biology have underscored the remarkable plasticity of pluripotent stem cells (PSCs), with their ability to self-organize and differentiate, can be induced to form embryo-like structures (including blastoids, embryoids, and gastruloids) under specific conditions, providing a valuable tool for in vitro investigation of embryonic development^1-4^.

Blastoids are advanced models for studying early embryo development in vitro, initially developed in mice by assembling embryonic stem cells (ESCs) or extended pluripotent stem cells (EPSCs) with trophoblast stem cells (TSCs), or through EPSC differentiation and self-organization, have also been successfully generated in humans, bovines, and monkeys^5-10^. In mouse and bovine models, both embryonic and extraembryonic stem cells are necessary for assembling embryo-like structures^5,11^. In contrast, human and monkey naive embryonic stem cells (hnESCs) can be directly induced into blastoids under specific induction protocols, indicating a robust model for studying human blastocyst development in vitro^3,9,11-13^. This might be due to the higher plasticity of human and monkey naive embryonic stem cells. Pig blastocyst-like structures have been induced from porcine ESCs using the suspension EB method, but the efficiency of obtaining blastocyst-like structures with this method is less than 20%^14^. However, the reported efficiency of blastocyst-like structure formation in humans, cattle, and monkeys is above 70%^10,15,16^. In comparison, the current efficiency of inducing porcine blastocyst-like structures is too low.

Porcine Expanded Potential Stem Cells (pEPSCs) have been established, with their ability to possess both embryonic and extra-embryonic differentiation potential under certain conditions^17^. The differentiation potential of pEPSCs allows us to establish an efficient porcine blastoid induction system. In this study, based on an optimized protocol, we generated porcine blastoids using porcine EPSCs with an efficiency of over 80%. Extensive morphological, molecular, and functional characterization revealed that these blastoids could mimic the fundamental blastocyst-like structure.

## Results

To establish a condition for inducing pig blastocysts, we seeded 0.5 × 10^4^/cm^2^ porcine EPSCs in a plate coated with 0.5% Matrigel. The WNT, FGF, and STAT3 signaling pathways indeed play important roles in the lineage differentiation and pluripotency regulation of embryonic stem cells^18-21^. Then we conducted a screening of cytokines and small molecules associated with the WNT, Activin/Nodal, STAT3, and FGF signaling pathways (Fig. 1A). The WNT signaling pathway plays a crucial role in the maintenance and differentiation of embryonic stem cells, promoting self-renewal and pluripotency while also guiding lineage specification during development^22,23^. Additionally, it has been reported that excessive WNT signaling can lead to embryonic stem cell differentiation. To obtain differentiated cells, we attempted to adjust the concentration of CHIR99021; however, we did not achieve significant differentiation of the cells (Fig. 1B). FGF can bias the fate of inner cell mass (ICM), likely acting through the RAS-RAF-MEK-ERK/MAPK pathway, and high levels FGF direct ICM cells toward the hypoblast (HYPO)^24-26^. We attempted to add FGF2 to the pEPSCs medium (Fig. 1B). However, we did not observe the appearance of Hypo-like cells. LIF, IL6, and sIL6R are activators of the STAT3 signaling pathway, and they play important roles in the culture of embryonic stem cells^27-29^. Recent studies have shown that IL6 plays an important role in human Hypo lineage differentiation^30^. Therefore, we added them to the pEPSCs medium. However, after adding IL6, sIL6R, and FGF2, we did not observe significant changes. XAV939 is a potent Tankyrase inhibitor that stabilizes AXIN protein and inhibits the Wnt/β-catenin signaling pathway, plays a significant role in maintaining embryonic stem cell pluripotency and inhibiting lineage differentiation by modulating the Wnt/β-catenin signaling pathway^31^. In particular, Src blockade partly arrested morula and affected trophectoderm and primitive endoderm (PrE)^32,33^. It is possible that components XAV939 and WH4023 in the pEPSCs medium strongly inhibit differentiation. Upon removing XAV939 and WH4023 from the pEPSCs medium, we observed partial differentiation in some clones, but did not observe trophoblast-like cells (Fig. 1B). Minocycline hydrochloride (MiH) is a HIF-1α inhibitor, Adding MiH to mouse embryonic stem cells (mESC) can induce them to transition into cells resembling trophoblast stem cells (TSC)^34^. The addition of Minocycline hydrochloride in the absence of XAV939 and WH4023 creates a unique environment that promotes the differentiation of pEPSCs into TSC-like cells (Fig. 1B). This suggests that Minocycline hydrochloride may play a role in promoting trophoblast differentiation by modulating specific pathways or factors that drive this process. When pEPSCs are cultured in medium 12, we frequently observe the formation of suspended cyst structures in cultures by day 2 post-passaging (Fig. 1B). By day 4, these structures increase in size (78.6 ± 10.6 μm), and some of them develop into blastocyst-like structures (Fig. 1B-F). However, this method results in a blastocyst-like structure formation efficiency of less than 20% (Fig. 1E).

**Figure 1.**
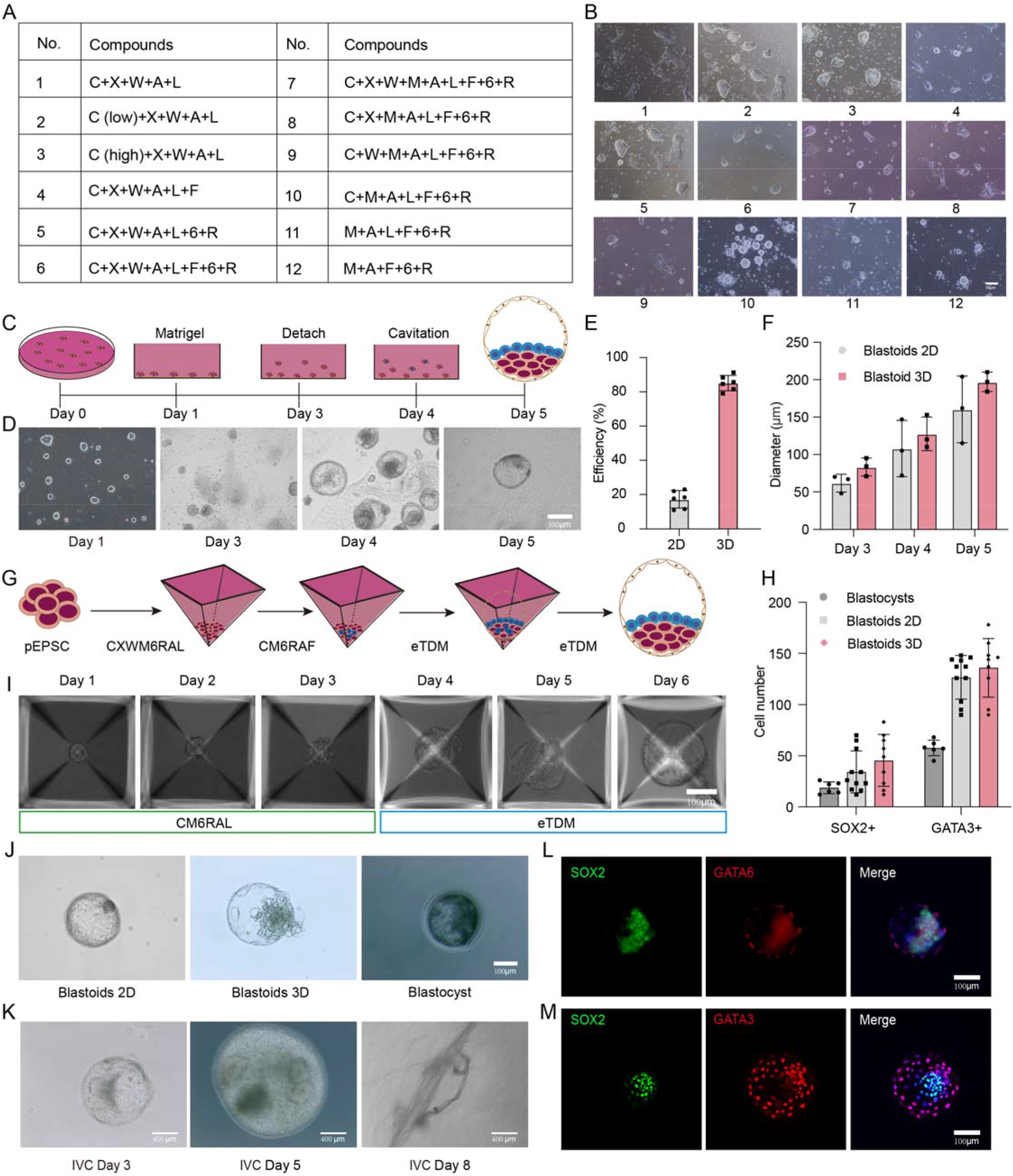
Generation of pig blastoids from pEPSCs. A. Small molecules used in pig blastoid induction and their targets, CHIR99021 (C), XAV939 (X), Minocycline hydrochloride (M), WH4023 (W), Activin A (A), LIF (L), FGF2 (F), IL-6 (6), sIL-6R (R). B. Morphological of pEPSCs after treatment of different chemical conditions. Scale bar, 50 μm. Representative of three independent experiments. C. Schematic diagram of the generation of porcine blastocysts in a 2D culture. D. Morphology of the generation of porcine blastocysts in a 2D culture. Scale bar, 100 μm. E. Blastoid formation efficiency of pig blastoids medium12 to mTDM condition (n = 6). F. Mean diameter of pig blastoids induced under 2D and medium12 to mTDM condition (n = 5) and IVF pig blastocyst (n = 5). G. Schematic diagram of the generation of porcine blastocysts in a 2D culture. H. Total cell numbers of per pig blastoids from 2D and 3D (n = 5 biological replicates) and blastocyst (n = 5 biological replicates). I. Morphology of the generation of porcine blastocysts in a 3D system. Scale bar, 100 μm. J. Representative images of pig blastoids from 2D (left) and 3D (middle) system on day 6 and pig E6 blastocyst (right). Scale bars, 100 μm. K. The dynamic of the morphology of pig blastoid IVC from day 1 to day 3 and day 8. Scale bars, 400 μm. L. Representative immunofluorescent staining of markers of epiblast marker (SOX2), hypoblast marker (GATA6) in reconstructed pig blastoids on day 6. Scale bars, 100 μm. M. Representative immunofluorescent staining of markers of epiblast (SOX2) and trophectoderm (GATA3) in reconstructed pig blastoids from 3D condition on day 6. Scale bars, 100 μm.

To efficiently generate pig blastoids from pEPSCs, we transferred the pEPSCs to aggrewell for 3D induction. Continuously using medium 12 did not induce blastocyst formation in the 3D culture. Trophoblast differentiation medium (TDM) is an essential culture system for generating blastocyst-like structures from human and monkey cells. We switched to TDM at different time points during induction with Medium 12 (Fig. G). This optimized step supported the formation of pig blastoids with high efficiency (80% ± 10%) within 6 days (Fig. E-J). Immunofluorescence staining showed that the blastocyst-like structures were usually composed of inner EPI-like cells (SOX2+), Hypo-like cells (GATA6+) and outer TE-like cells (GATA3+) (Fig. 1J, L, M). Next, we evaluated the in vitro growth of pig blastoids under a 3D suspension culture system to investigate their post-implantation developmental potential. Blastoids at day 6 with morphology of apparent cavity and ICM-like cells inside were chosen for prolonged culture. The expanded pig blastoids were transferred from an aggrewell plate to an 8-well dish and cultured in vitro prolonged culture medium 1 (IVC1). Many of the pig blastoids attached to the dish after 24 h, at which point the medium was changed to IVC2 and cycled every other day. We observed that the porcine blastoids cultured in vitro gradually increased in size, reaching approximately 600 μm by day 3. Additionally, When the blastocyst-like structures were cultured in IVC2 medium until day 5, the diameter of the porcine blastocyst-like structures grew to approximately 1400 µm, and the elongation of trophoblast cells was observed on day 8 (Fig. 1K).

To determine the transcriptional states of pig blastoid cells, we performed single-cell RNA sequencing (scRNA-seq) using the 10 x Genomics Chromium platform and carried out integrated analysis with Smart-seq2 single-cell transcriptomes derived from multiple crossbred Large White and Landrace sows (2–3 years old) between days 4 and 11 after artificial insemination pig embryos^29^. Joint uniform manifold approximation and projection (UMAP) embedding revealed pig blastoid-derived cells clustered with blastocyst-derived cells (Fig. 2A, D, E, and F). To further evaluate the temporal identity of blastoid cells, we performed two pseudo bulk analyses on the 10 x blastoid data at a low and a high cluster resolution, to compensate for the differences in sequencing depth to Smart-seq2 data. We found that different embryo datasets were orderly arranged on the principal component analysis (PCA) plot according to their developmental time, and blastoid cells were mapped closer to blastocyst cells (Fig. 2G). We annotated the eight identified cell clusters based on marker gene expression and overlap with cells from bovine embryos (Fig. 2B, D, E, F, G and H). Cluster 7 expresses EPI markers, e.g., POU5F1, SOX2 and LIN28A, and is annotated as ELCs; cluster 1 expresses HYPO markers, e.g., SOX17, GATA4 and FOXA2, and thus represents HLCs; Cluster 4 expresses TE markers, e.g., GATA2 and GATA3, and is annotated as TLCs; cluster 0, cluster 2 and cluster 6 is mostly composed of cells from pre-blastocyst stage embryos (named pre-lineage). These analyses confirm that pig blastoids recapitulate major pig blastocyst cell types at the transcriptomic level.

**Figure 2.**
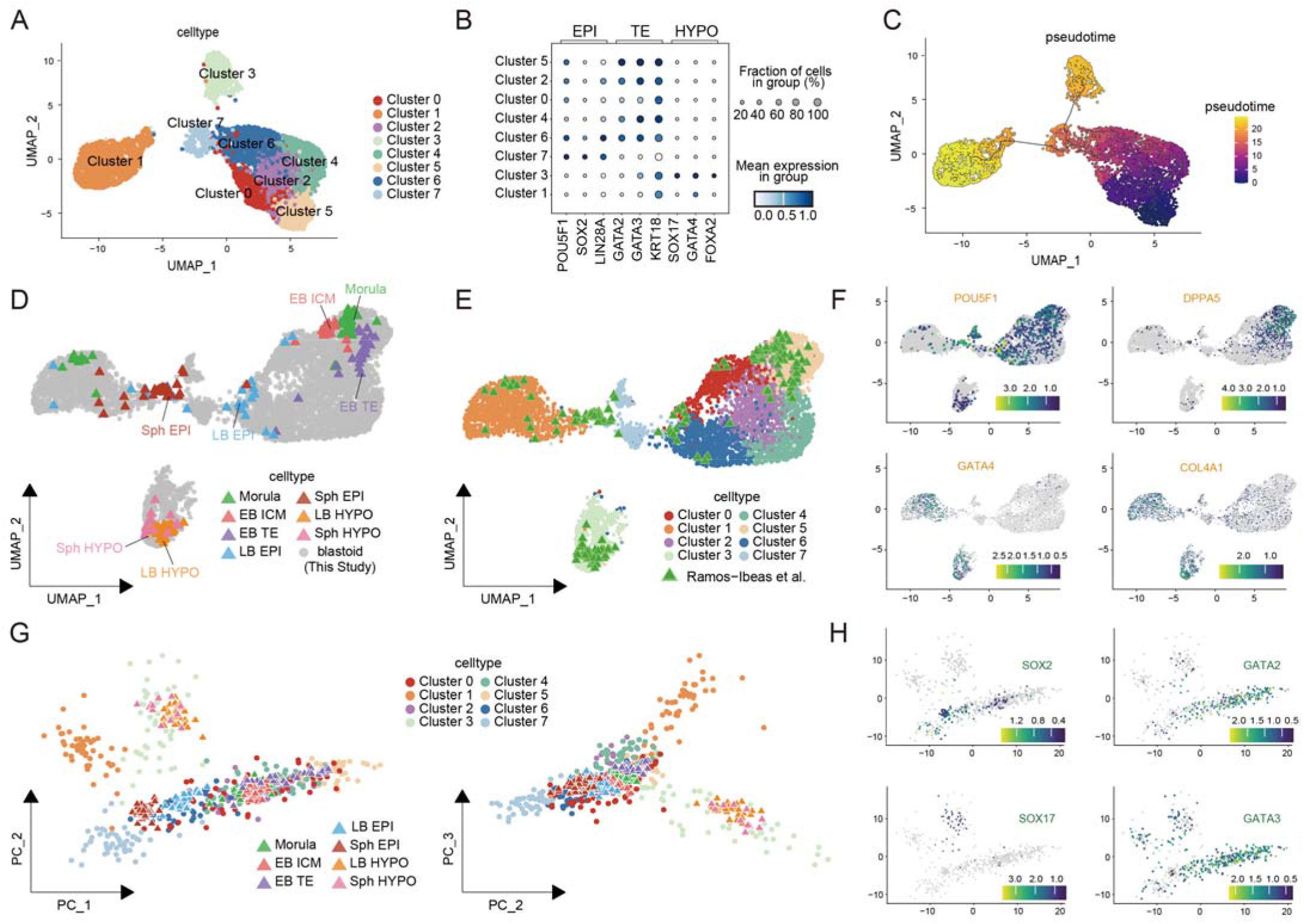
Single-cell transcriptome analysis of pig blastoids. A. Uniform Manifold Approximation and Projection (UMAP) of blastoids cell populations based on RNA expression. N=5827 total cells. B. Dot plot indicating the expression of marker genes in the trophectoderm (TE), hypoblast (HYPO), and epiblast (EPI) during pig embryonic development. C. RNA velocity pseudotime analysis overlaid on UMAP depicting the cell trajectories. D. Joint UMAP of cells from blastoids and blastocyst stage embryos (Ramos-Ibeas et al.), points colored according to blastocyst. cell type and blastoid cells (grey). E. Joint UMAP of showing the integrating result of cells from blastoids and blastocysts data (Ramos-Ibeas et al.). Color represents the subtype annotation. F. The UMAP visualization displaying the expression of key markers for TE, HYPO, and EPI: POU5F1, DPPA5, GATA4, and COL4A1, correspondingly. G. Principal component analysis (PCA) plot of cell types from blastoids and blastocysts data (Ramos-Ibeas et al.). PCA1 is represented on the X-axis, while PCA2 is depicted on the Y-axis (left). PCA2 is shown on the X-axis, and PCA3 is displayed on the Y-axis (right). H. PCA displaying the expression of classical markers for TE, HYPO, and EPI: SOX2, SOX17, GATA2, and GATA3, respectively.

## Discussion

Our study shows that optimizing the culture medium and conditions can significantly enhance the efficiency of pig blastocyst-like formation by continuing to refine and optimize this protocol, we aim to unlock new possibilities in the field of animal reproduction and contribute to the improvement of livestock breeding practices. In human, bovine and mouse systems, the role of FGF signaling in promoting the formation of hypoblast-like cells from the inner cell mass (ICM) is well-documented^35-37^. Similarly, in our porcine system, we found that FGF2 is critical for directing ICM cells toward a hypoblast fate, likely via the RAS-RAF-MEK-ERK/MAPK pathway. However, the presence of inhibitors like XAV939 and WH4023 in the pEPSCs medium initially hindered differentiation, a challenge also reported in human blastoid systems^13,17^. In mouse blastoids, the use of Minocycline hydrochloride (MiH) to induce trophoblast stem cell-like properties highlights the significance of HIF-1α inhibition in trophoblast differentiation^34^. Then, we added MiH and observed the appearance of TE-like cells, while also maintaining the clones of EPSCs. In this 2D induction system, we obtained porcine blastoids with an efficiency of less than 20%. This efficiency is similar to the latest reported efficiency for porcine blastocyst induction^14^.

The switch to a 3D culture system and the incorporation of Trophoblast Differentiation Medium (TDM) significantly improved the efficiency of blastoid formation in our porcine model, achieving approximately 80% efficiency within six days. This efficiency is notably higher than that achieved with Medium 12 alone and aligns with efficiencies reported in human, pig and monkey blastoid systems using similar 3D culture approaches^7,10,14,38^.

The transcriptional profiles of porcine blastoids closely resemble those of natural blastocyst-derived cells, as revealed by single-cell RNA sequencing (scRNA-seq). This resemblance is consistent with findings in human, bovine, and mouse studies, where blastoid cells exhibit gene expression patterns similar to their in vivo counterparts^29^. The clustering of blastoid-derived cells with natural blastocyst cells in UMAP analysis further supports the fidelity of our induced blastoids to natural embryonic development. In bovine systems, the challenge of achieving high efficiency in blastoid formation is also observed, with studies often reporting lower efficiencies compared to human and mouse models. Our study’s success in achieving high efficiency in porcine blastoids underscores the potential of optimizing culture conditions and modulating specific signaling pathways to overcome species-specific barriers.

Our study demonstrates that optimizing culture conditions and precisely modulating signaling pathways are crucial for the efficient induction of porcine blastoids from pEPSCs. The high efficiency and fidelity of the generated blastoids provide a valuable model for studying early embryogenesis in pigs and offer promising applications in regenerative medicine and reproductive biology.

## Method details

### In vitro fertilization

Porcine ovaries were harvested and placed in saline solution containing penicillin and streptomycin at 32–37°C. Antral follicles (2–8 mm in diameter) were aspirated using 10-gauge needles. High-quality oocytes, characterized by uniformly granulated cytoplasm and at least three layers of compact cumulus cells, were meticulously chosen. These oocytes were then cultured in maturation medium (TCM-199 supplemented with 0.07 mg/ml cysteine, 0.05 mg/ml epidermal growth factor (EGF), 0.5 mg/ml luteinizing hormone (LH), and 0.5 mg/ml follicle-stimulating hormone (FSH)) under conditions of 38.5 °C and a 5% CO_2_ atmosphere. After 42 hours of maturation, mature porcine oocytes exhibiting the first polar body were identified and selected for subsequent experiments. Each group of 30 mature oocytes was individually placed in a 50 μl droplet of modified Tris-buffered medium (mTBM) for fertilization. Fresh sperm, initially stored at 17 °C, was briefly equilibrated to 38.5 °C before being washed twice in Dulbecco’s phosphate-buffered saline (DPBS, Sigma) supplemented with 0.1% (w/v) bovine serum albumin (BSA, Sigma). The washed sperm suspension was then added to the droplets containing oocytes at a final concentration of 3×10^^5^ cells /ml. The oocyte-sperm mixtures were subsequently incubated at 38.5 °C for 6 hours. Finally, the fertilized embryos were cultured in porcine zygote medium-3 (PZM-3) in a 4-well plate, maintained at 38.5 °C under a 5% CO_2_ atmosphere.

### Establishment of Porcine EPSCs stem cell culture

We established porcine expanded pluripotent stem cells (pEPSCs) from porcine IVF blastocysts by referencing the method developed by Pentao Liu^17^. Pig IVF blastocysts from day 5 were used to obtain ICMs by placing them in Ca^2+^-TL-HEPES medium by microsurgery using ophthalmic scissors. Isolated ICMs were cultured for 7 d on MEF cells in pEPSCs medium, in the presence of 10 μm Y27632, until initial outgrowths could appear. The outgrowths were mechanically isolated and reseeded onto fresh STO cells in pEPSCs medium. The cells formed well-defined porcine EPSCs colonies 3 d later EPSCs cells were maintained on MEF feeder layers and enzymatically passaged every 3–5 d by a brief PBS wash followed by treatment with 0.25% trypsin/EDTA for 3–5 min. The cells were dissociated and centrifuged (300g for 5 min) in 10% fetal bovine serum (FBS)-containing medium. After removing the supernatant, the porcine/human EPSCs were resuspended and seeded in pEPSCs supplemented with 5 μm Y27632 (Tocris, cat. no. 1254). The addition of 5% FBS (Gibco, cat. no. 10270) and 10% KnockOut Serum Replacement (KSR; Gibco, cat. no. 10828-028) improved the survival of cells during passaging. The medium was switched to pEPSCs only 12–24 h later. pEPSCs cell N2B27 based media. N2B27 basal media (500 ml) was prepared as follows: 482.5 ml DMEM/F-12 (Gibco, cat. no. 21331-020), 2.5 ml N2 supplement (Thermo Fisher Scientific, cat. no. 17502048), 5.0 ml B27 supplement (Thermo Fisher Scientific, cat. no. 17504044), 5.0 ml 100× glutamine penicillin-streptomycin (Thermo Fisher Scientific, cat. no. 11140-050), 5.0 ml 100× NEAA (Thermo Fisher Scientific, cat. no. 10378-016) and 0.1 mM 2-mercaptoethanol (Sigma, cat. no. M6250). To make pEPSCs (500 ml), the following small molecules and cytokines were added into 500 ml N2B27 basal media: 0.2 μm CHIR99021 (GSK3i; Tocris, cat. no. 4423), 0.3 μm WH-4-023 (Tocris, cat. no. 5413), 2 μm Minocycline hydrochloride (Santa Cruz Biotech-nology, Cat. No. SC-203339), 2.5 μm XAV939 (Sigma, cat. no. X3004) or 2.0 μm IWR-1 (Tocris, cat. no. 3532), 65.0 μg ml^−1^ vitamin C (Sigma, cat. no. 49752-100G), 10.0 ng ml^−1^ LIF (Stem Cell Institute (SCI), University of Cambridge), 20.0 ng ml^−1^ Activin (SCI), 10.0 ng ml^−1^ FGF2 (Peprotech, cat. no. 100-18B), 10.0 ng ml^−1^ IL-6 (Peprotech, cat. no. 200-06-100 ug), 10.0 ng ml^−1^ sIL-6R (Peprotech, cat. no. 100-18B) and 0.3% FBS (Gibco, cat. no. 10270). All cell cultures in this paper were maintained at 37 °C with 5% CO_2_, unless stated otherwise.

### Blastoid formation

For blastoid formation, pEPSCs single cells were collected as stated above. pig EPSCs were washed with 1x PBS, dissociated with Trypsin for 5 minutes at 37 °C, with constant pipetting every 2-3 minutes and inactivated with DMEM-F12 containing 10% fetal bovine serum (FBS). Cells were washed twice and on final resuspension in their normal culture media with 1 x CEPT. To deplete iMEF cells, collected cells were placed in precoated 12 well plates (Corning) with 0.1% gelatin and incubated for 45 minutes at 37 °C. Single-cell dissociation was made by gentle but constant pipetting and by passing the cells through a glass capillary pulled to an inner diameter of 50-100 μm (micropipette puller, Sutter Instruments), hermetically attached to a p200 pipette tip. After single-cell dissociation, cells were collected and strained using a 70 μm (EPSCs) (Corning). This same single-cell dissociation procedure was used for blastoids processing for 10 x genomics. Cells were stained with 1x trypan blue and manually counted in a Neubauer chamber. Current protocol is optimized for 20 pEPSCs per well in a 1200 well Aggrewell 400 microwell culture plate (Stemcell technologies) for 24,000 of each cell type per well. Each well was precoated with 500 ml of Anti-Adherence Rinsing Solution (Stemcell technologies) and spun for 5 minutes at 1500 rcf. Wells were rinsed with 1 ml of PBS just before aggregation. An appropriate number of cells for the wells to be aggregated was centrifuged at 100 x g for 2 minutes and resuspended in 1ml of medium12 (N2B27 base, 1% BSA, 0.5 x ITS-X, 20 ng/mL IL-6, 20 ng/mL sIL6R, 10 ng/ml LIF, 10 ng/ml FGF2, 1 μm Minocycline hydrochloride, 1 μm CHIR99021) per well, supplemented with 1 x CEPT. To ensure even distribution of the cells within each microwell, cells were gently mixed by pipetting with a P200 pipette, then the plate was centrifuged at 100 x g for 2 minutes and put in a humidified incubator at 37 with 5% CO_2_ and 5% Oxygen (NuAire). It is important to have viable, MEF free, cell debris free, and evenly distributed cells as any of these factors can negatively affect blastoid formation.

### Immunofluorescent staining

Samples (Cells, blastoids and blastocysts) were fixed with 4% paraformaldehyde (PFA) in DPBS for 20 min at room temperature, washed three times with DPBST (DPBS,0.1% Triton X-100) for 15 minutes and permeabilized with 0.5% Triton X-100 in DPBS for 30 minutes. Samples were then blocked with blocking buffer (DPBST containing 3% BSA) at room temperature for 1 h, or overnight at 4°C. To facilitate blastoids staining, blastoids were gently washed out of the aggrewell plate and then transferred in a 4 well plate (Thermo Fisher) containing wash buffer with mouth pipette and then blastoids were moved from one well to another between steps. Primary antibodies were diluted in blocking buffer according to supplementary table 1. Blastoids were incubated in primary antibodies in 4 wells overnight at 4°C.Samples were washed three times for 15 minutes with wash buffer and incubated with fluorescent-dye conjugated secondary antibodies (AF-488, AF-594 or AF-647, Invitrogen) diluted in blocking buffer (1:200 dilution) for 1 h at room temperature and then washed three times with DPBST. Finally, cells were counterstained with 300 nm. 4′,6-diamidino-2-phenylindole (DAPI) solution at room temperature for 5 min.

### Imaging

Phase contrast images were taken using a hybrid microscope (ZEISS axio observer z1) equipped with objective x2/0.06 numerical aperture (NA) air, x4/0.13 NA air, x10/0.7 NA air and 20x/0.05 NA air. Fluorescence imaging was performed on 8 well u-siles (Ibidi) on a Nikon A1 spinning-disk super resolution by optical pixel reassignment (SoRa) confocal microscope with objectives x4/0.13 NA, a working distance (WD) of 17.1nm, air, ×20/0.45 NA, WD 8.9-6.9 nm, air; ×40/0.6 NA, WD 3.6-2.85 nm, air.

### Imaging analysis

The imaging experiments were replicated at least twice and yielded consistent outcomes. In the figure captions, ‘n’ indicates the number of biological replicates. Initially, raw images underwent processing in Fiji to generate maximal intensity projections (MIP) and to export representative images. Subsequently, nuclear segmentation was conducted using Ilastik. MIP images and segmentation masks were then analyzed in MATLAB (R2022a) using custom code, accessible in a public repository. For each cell within every field, the nuclear-localized fluorescence intensity was quantified and normalized to the corresponding DAPI intensity. These intensity values for all cells were graphed as mean ± standard deviation. Total cell counts and the numbers of SOX2, and GATA3-positive cells were calculated using Imaris (v.9.9, Oxford).

### In vitro growth

Porcine blastoids were manually isolated using a mouth pipette, washed with PBS and transferred into 24-well plate pre-coated with 0.5% matrigel containing IVC-1 medium. The medium was switched to IVC-2 medium 24hrs later on which time almost all blastoids attached and the medium was changed every 2 days. IVC-1 and IVC-2 media were prepared using the following: IVC-1: DMEM/F12 (Gibco) supplemented with 20% (v/v) FBS, 2 mM L-glutamine (Gibco), 1 x ITS-X, 8 nm. β-oestradiol (Sigma), 200 ng/mL progesterone (Sigma), 25 μm N-acetyl-L-cysteine (Sigma); IVC-2: DMEM/F12 supplemented with 30% (v/v) KSR, 2 mM L-glutamine, 1 x ITS-X, 8 nm. β-oestradiol, 200 ng/mL progesterone, 25 μm N-acetyl-L-cysteine.

### Single cell library construction

Porcine blastoids were single cell dissociated and strained cells were prepared as stated adobe. Cells were washed in PBS containing 0.04% BSA and centrifuged at 300g for 5 min. Cell were resuspended in PBS containing 0.04% BSA at a single cell suspension of 1,000 cells/µL. Cells were loaded into a 10 x Genomics Chromium Chip following manufacturer instruction (10x Genomics, Pleasanton, CA, Chromium Next GEM Single Cell GEM, Library & Gel Bead Kit v3.1) and sequenced by Illumina NextSeq 500/550 sequencing systems (Illumina).

### Preparing single-cell data for analysis

The study utilized the Cell Ranger pipeline version 7.2.0 with default parameters to analyze single-cell data obtained from 10 x Genomics. The expression count matrix was generated using standard settings. The reference genome and gene annotation information for pigs (Sscrofa10.2) were sourced from Ensembl (https://ftp.ensembl.org/pub/release-87/) and prepared using Cell Ranger’s mkgtf and mkref tools with their default settings.

The Seurat v4 (version 4.3.0) software package was used to control the single-cell quality. Cells were subjected to filtration based on specific parameters to minimize the presence of multiples and non-viable cells. Single cells with the number of detected genes (nFeature_RNA) above 2,000, detected transcripts (nCount_RNA) above 5,000 and mitochondrial RNA gene counts below 15 percent were retained to next analysis.

The Smart-seq2 data and its corresponding annotation information were acquired from the Gene Expression Omnibus (GEO) database with the accession number GSE112380. Subsequently, the R package biomaRt (version 2.58.0) was employed to convert the Ensembl IDs to gene symbols. In order to minimize differences caused by data processing, the useMart function was utilized with the same genome reference (Sscrofa10.2).

### Data normalization and clustering

The log-percentage value was utilized in the NormalizeData function to standardize the expression matrix of individual cells, thus reducing potential distortions in gene expression levels caused by variations in sequencing depths and methods. The PCA analysis was carried out using the RunPCA function on a set of 2000 variable genes identified through the FindVariableFeatures function. Subsequently, the UMAP dimensionality reduction was performed using the RunUMAP function, based on the clusters identified by the FindClusters function.

### Data integration

The FindIntegrationAnchors function from the Seurat package was employed with a specified parameter (k.anchor = 10) to identify common anchors between two datasets. Subsequent to this step, the data integration process was carried out using the IntegrateData function based on the identified anchors. After integration, the ScaleData and RunPCA functions were applied to the merged dataset. Furthermore, the cell distribution was visualized using the RunUMAP function with non-default parameter: ‘dims = 1:20’.

### Principal component analysis

Based on the significantly larger number of cells in this study compared to the Smart-seq2 data, before integration, a subset of 50 cells from each cell type was randomly selected to create a new dataset. This dataset was then combined with the reference data using the specified data integration method with default parameters. Subsequently, the principal component analysis (PCA) data generated by the RunPCA function, which used the top 1000 features identified by the VariableFeatures function.

### Pseudotime analysis

The R package monocle3 (version 1.3.5) was used for calculating the pseudotime of single cells. The monocle3 object was formed using the new_cell_data_set function with the count matrix from the Seurat object. The monocle3 object was processed using the preprocess_cds function, incorporating the UMAP reduction information from Seurat. Subsequently, the pseudotime trajectory was established through the learn_graph and order_cells functions. Finally, the visualization of the pseudotime trajectory was achieved using the plot_cells function.

## Author contributions

Y.W., S.Z. conceptualized the idea, Y.W., Z.L. designed, analyzed, and interpreted the experimental results. Z.L. performed blastoid generation and extended invitro culture experiments with the help of S.Y.. J.Z., N.H. J.Z. helped with immunostaining. X.D. and X.G. provided with in vitro fertilization of pig embryos. L.Z. prepared scRNA-seq library. X.Q. performed scRNA-seq analysis. Y.W. and S.Z. supervised the study. Z.L. and Y.W. wrote the manuscript with inputs from all authors.

## Acknowledgements

We thank Dr. Yiliang Miao from Huazhong agriculture university for providing pig EPSC cells. Y.W. is funded by STI 2030-Major Projects (2023ZD0407504), The Innovative Project of of State Key Laboratory of Animal Biotech Breeding (2023SKLAB1-2); China Agricultural University Young Talent Program in Life Science (Grant No. 001). S.Z. is funded by the National Key R&D Program of China, Grant No. 2021YFD1200301.

